# Gain efficiency with streamlined and automated data processing: Examples from high-throughput monoclonal antibody production

**DOI:** 10.1101/2023.12.14.571214

**Authors:** Malwina Kotowicz, Sven Fengler, Birgit Kurkowsky, Anja Meyer-Berhorn, Elisa Moretti, Josephine Blersch, Magdalena Shumanska, Gisela Schmidt, Jakob Kreye, Scott van Hoof, Elisa Sánchez-Sendín, S. Momsen Reincke, Lars Krüger, Harald Prüss, Philip Denner, Eugenio Fava, Dominik Stappert

## Abstract

Data management and sample tracking in complex biological workflows are essential steps to ensure necessary documentation and guarantee the reusability of data and metadata. Currently, these steps pose challenges related to correct annotation and labeling, error detection, and safeguarding the quality of documentation. With growing acquisition of biological data and the expanding automatization of laboratory workflows, manual processing of samples is no longer favorable, as it is time- and resource-consuming, is prone to biases and errors, and lacks scalability and standardization. Thus, managing heterogeneous biological data calls for efficient and tailored systems, especially in laboratories run by biologists with limited computational expertise. Here, we showcase how to meet these challenges with a modular pipeline for data processing, facilitating the complex production of monoclonal antibodies from single B-cells. We present best practices for development of data processing pipelines concerned with extensive acquisition of biological data that undergoes continuous manipulation and analysis. Moreover, we assess the versatility of proposed design principles through a proof-of-concept data processing pipeline for automated induced pluripotent stem cell culture and differentiation. We show that our approach streamlines data management operations, speeds up experimental cycles and leads to enhanced reproducibility. Finally, adhering to the presented guidelines will promote compliance with FAIR principles upon publishing.

## 1. Introduction

Over the last few decades, technological advancements in fields such as imaging, laboratory automation, computing, and data analysis have revolutionized the way biologists work and handle data (1–4). High-throughput (HT) and high-content (HC) studies are no longer exclusive to large, specialized labs but are gaining popularity in research conducted by smaller, independent teams (4–6). This trend is expected to continue, as smaller biology labs increasingly adopt HT/HC techniques due to decreasing costs, thereby generating large amounts of biological data. Additionally, many funding agencies and scientific journals have emphasized the value of HT/HC techniques in producing reliable and comprehensive data, which will further incentivize individual groups to incorporate HT/HC methods to generate high-quality research and stay competitive (7–10).

Biological data is heterogeneous by nature and often includes experimental readouts, curated annotations, and metadata, among other type of data. The increasing size and complexity of biological datasets call for effective means to manage data in its complete life cycle throughout a workflow, and the generation, processing, analysis, and management of heterogeneous biological data require tailored systems to improve data governance (11–13).

The increasing use of laboratory automation and the generation of experimental workflows with complex structures pose a unique challenge in backtracking and identifying samples and their related metadata. Each step of the workflow impacts the final data, and if problems arise, it can be difficult to backtrack and pinpoint where errors occurred. Likewise, the reproducibility of complex biological workflows is closely tied to precise record-keeping, especially as new techniques are introduced. As wet lab experiments are often complex, time-sensitive, and involve many researchers, the quality of documentation can be compromised. Moreover, manual data curation is time-consuming, labor-intensive, prone to human error, and at risk of biases, as it relies on the individual expertise of the curator. Any error in data curation compromises data integrity and can lead to incorrect conclusions, inefficient workflows, and the inability to reuse the data. Similarly, manual integration of data from multiple sources lacks standardization, has limited scalability, and can hinder the early detection of errors (14).

To ensure data integrity in workflows and prevent potential data loss, strict quality control measures and careful monitoring of workflow steps are necessary. Although many systems exist for managing large datasets in biology (15), they are mainly implemented in larger, specialized facilities with teams trained in computer science. In smaller, individual labs, dedicated informatics staff may not be available and biologists are required to learn complex tools and technologies for data processing, despite lacking prior experience and facing severe time and resource constraints. Overall, there is an urgent need for design guidance for data processing solutions in biology workflows.

Here, we present the recently established pipeline for modular data processing that facilitates and documents the complex production of monoclonal antibodies (mABs) derived from individual B-cells. Implementing our data management system reduced the time spent on data processing by over one-third and improved data reliability. Our strategy proves that, with moderate effort, biologists can set up an efficient, rewarding, systematic approach to routine data processing tasks. This approach will: i) simplify documentation; ii) facilitate reproducibility and improve accuracy by eliminating errors related to manual data handling; iii) speed up data processing, accelerating the generation of reliable insights and freeing hands for other tasks; iv) standardize data processing procedures, enabling comparison of results across series of experiments or labs; and v) support scalability, as modules of data processing pipelines can be up- or down-scaled to handle varied data amounts, adapting to changing research needs. Finally, and to demonstrate the versatility and transferability of our approach, we apply it to the development of a data processing pipeline for automated stem cell culture. We show that our design guidelines can serve as best practice recommendations for other biologists and be a step towards greater reproducibility, efficiency, and standardization of workflows in biology.

### Box 1.

**Terminology**

This box provides definitions for selected terms throughout the text.

**Data curation**

the process of cleaning, organizing and standardizing data towards greater quality, utility and long-term preservation. Data curation is part of data processing.

**Data processing**

all tasks that are performed on acquired data to prepare it for analysis, especially curation, formatting, transposition, joining, sub-setting and summarizing, but excluding data acquisition and storage. Data processing is a broad term, encompassing all data manipulation and transformation steps that facilitate extracting meaningful insights and, in the long term, knowledge.

**Data processing pipeline**

the sum of all data processing modules for one workflow, aimed at converting raw input data into usable information.

**Data repository**

storage space to catalog and archive data. Data is curated prior to deposit. Ideally, the storage space and contained data are managed by a database software that allows efficient retrieval of related data points.

**Knime module**

a workflow in Knime performing a series of related tasks or operations, where each operation is performed by a Knime “node” that represents the smallest operational unit of Knime.

**Metadata**

data that describes other, associated data. In the context of this work, metadata includes, but is not limited to: i) donor data associated with a sample (e.g., date of donation, cell number); ii) experimental settings and conditions (e.g., protocols, reagents, equipment); iii) association of experimental results with the original sample; iv) location of a given sample or its derivate.

**Module or data processing module**

an individual Python script or Knime workflow.

**Wet lab experiment or experimental procedure**

workflow steps that are conducted physically in the laboratory.

**Workflow**

systematic series of interconnected steps designed to achieve specific research objectives, including wet lab work, data processing and data storage.

## 2. Workflow for mAB production and data processing challenges

We have recently established a wet lab workflow that enables the production of over one thousand mABs per year from patient-derived single B-cells. An antibody is composed of two protein chains, the heavy (H) and the light (L) chain, where the light chain can be of either Kappa (κ) or Lambda (λ) type. Genetic information for both the H and L chains needs to be isolated and cloned. This enables *in vitro* generation of mABs with the same specificity as in the originating B-cell. The wet lab protocols have been outlined with minor deviations elsewhere (16–18). The procedures consist primarily of the following steps: cDNA is generated from individual B-cells. The cDNA serves as a template to amplify the H and L chains by PCR. As the L chain can be of κ or λ type, it is necessary to perform three parallel PCR reactions (see Figure S1). The PCR amplicons are analyzed for correct size by electrophoresis and sequenced upon positive evaluation. Primer pairs specific for the sequenced H and L chains, and equipped with overhangs for Gibson Assembly, are selected for a final PCR reaction (for a complete primer set, see Table S1 in Supplementary Material). The final PCR products, covering the specificity-determining variable regions of antibodies, are cloned by Gibson Assembly into plasmids encoding the constant regions of H or L chains. The assembled plasmids are amplified in bacteria, purified, and sequenced to confirm sequence identity, i.e., that no mutations causing failures on the protein level have been introduced (19). HEK cells are then transfected with pairs of plasmids encoding matching H and L chains. After transfection, mABs produced by HEK cells are secreted into the cell media, and their concentrations are measured.

Due to the complexity of the workflow, the production of mABs in a time- and resource-efficient manner implies a considerable data processing effort, as the following challenges must be addressed:

1. Wet lab experiments represent a selection funnel, wherein not all samples that enter the workflow proceed to the end. This is because they involve a series of steps that progressively narrow down the selection of samples, i.e., not all B-cells will give rise to functional antibodies. Optimization of resource usage, therefore, requires the selection of successful samples after each analysis step. The next wet lab step is only executed when a significant number of successful samples have accumulated after a set of wet lab experiments (e.g., one full plate of 96 samples).
2. A specific challenge to mAB production is their composition of H and L chains. As antibodies are composed of plasmid pairs, quality analysis requires the matching of heavy and light chains cloned from the same cDNA. Only if both heavy and light chains are positively evaluated are the chains pushed to the next step. If one chain of a pair is missing, several workflow steps must be repeated for the missing chain, increasing the complexity of both data processing and documentation.
3. Another challenge to mAB production is the variability of H and L chains derived from somatic recombination of V(D)J gene segments (20,21). The specificity-determining region of an H chain is composed of several V (variable), D (diversity), and J (joining) gene segments. The specificity-determining region of an L chain is similarly composed of several V (variable) and J (joining) gene segments. In order to clone H and L chains, a three-step PCR strategy is implemented (see Figure S1 and Table S1). First, forward and reverse primer mixes for each chain are used to amplify a given chain from cDNA. Then, a second PCR with primer mixes is performed to analyze the first amplicon by sequencing. Obtained sequences are analyzed for the specific V(D)J gene alleles, and based on this analysis, a specific primer pair is selected from a set of 69 primers to perform a final PCR step (19). Significant effort is required for processing the data, curating the data sets, and developing pipetting schemes for performing the final specific PCR.
4. The key to a time-efficient workflow is to shorten the data processing effort between two wet lab experiments. Due to the complexity of our workflow, data processing is time-consuming when done manually (≥33% of complete hands-on time for mABs production from cDNA). Analysis of samples, creation of pipetting schemes, and relating initial cDNA samples to experimental readouts are a challenge, as different machines use varying plastic ware layouts (e.g., 96- or 384-well plates column- or row-wise, individual culture plates or individual tubes).
5. Data generated on each mAB production step needs to be curated and documented before you can proceed to subsequent steps, analysis, troubleshooting, hypothesis formulation, and publication.

An organized and standardized approach to data processing and analysis can help address these and similar challenges in complex biological workflows. In this work, we propose a set of design guidelines for data processing pipelines, based on our experience with automated mAB production.

## 3. Getting started

When designing a workflow, it is crucial to conceptualize it in advance by defining the tasks, identifying their dependencies and contingencies, and determining data processing operations. This can be done in a few steps:

1. Start by outlining a series of step-by-step experiments (tasks) that constitute the workflow, specifying data to be retrieved, analyzed, and stored at each step. Determine the workflow endpoint to guide the design process. Visualizing the entire workflow helps to understand the experimental sequence and anticipate potential workflow expansion.
2. Establish methods to document samples’ metadata and experimental details, including protocols and data analysis procedures to ensure reproducibility and enable collaboration.
3. Consider the repetitions that may occur in case of experiment failure and how to handle them with respect to data storage.
4. Implement the ways to document the workflow runs and the results for quality control, error checking, data analysis and interpretation.

By defining these components, it becomes easier to identify the data processing operations that are required at each workflow step to form a functional pipeline (see Figure 1). In the next sections, we present design principles that guided the development of our data processing pipeline. We start with principles that give rise to functional pipelines, such as choosing the right technologies, applying modular design, ensuring interoperability between modules, and more. We cover the implementation of dedicated databases and further design guidelines that go beyond developing functional pipelines, but improve existing ones towards better efficiency, organization, and reproducibility.

**Figure 1.**
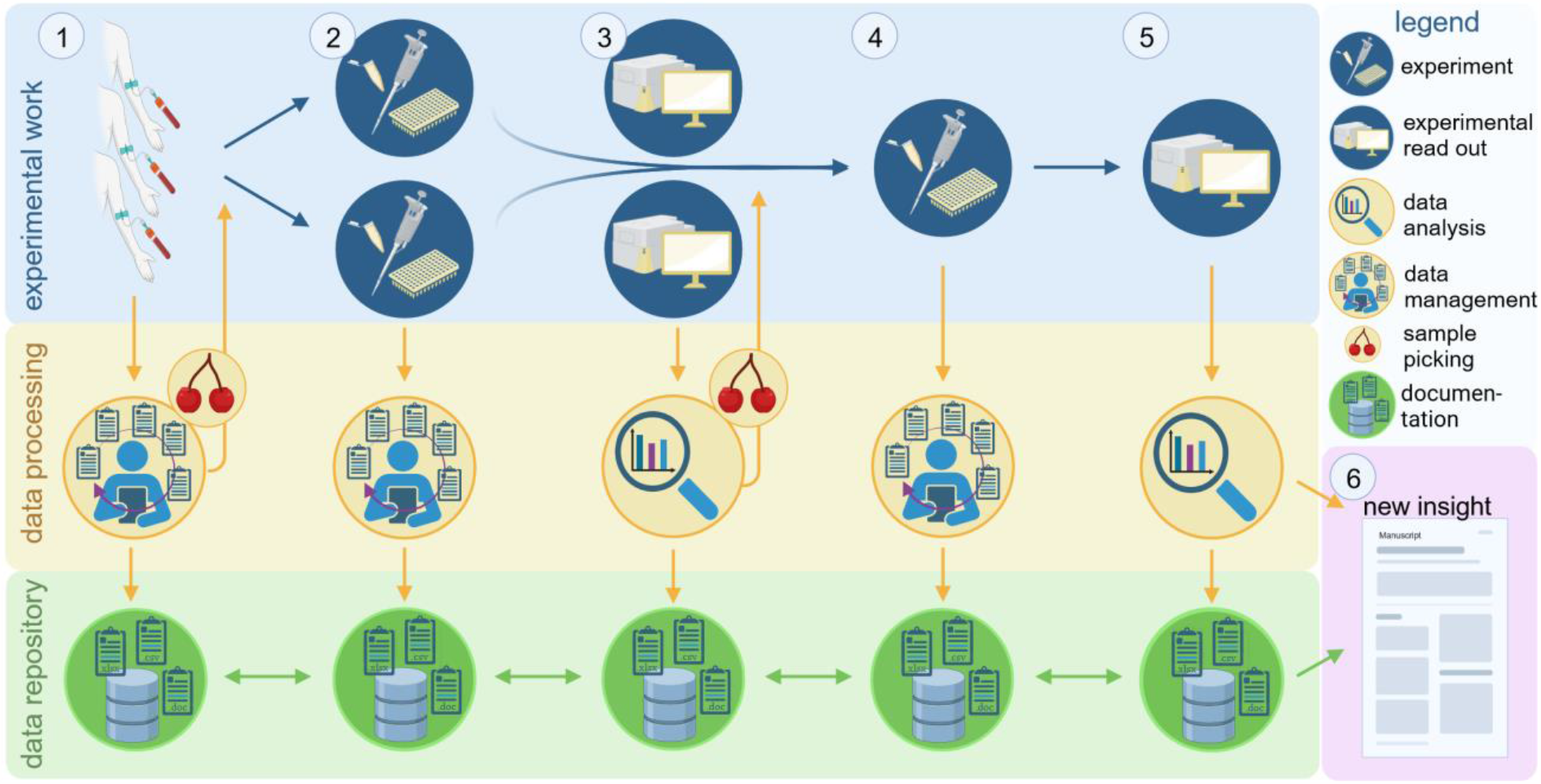
An example of a twostep wet lab workflow, demonstrating data processing needs. Data generated in wet lab experiments (experimental work, blue background, top) undergoes processing (data processing, yellow background, middle) for analysis and storage (data repository, green background, bottom). Biological samples and accompanying metadata are collected and must be curated and documented. Based on the metadata, subsets of samples are selected for analysis, such as those from wild-type or diseased subjects (1). The workflow starts with a wet lab experiment that is then documented (2) and analyzed (3). Next, the wet lab experiment is conducted on a subset of samples based on the analysis of the first experiment (4). The analysis of the second experiment closes the workflow cycle (5). Analyzing the results of the second experiment in the context of data (and metadata) gathered in the workflow cycle allows for supporting or refuting the hypotheses and yields novel insights (6).

When designing the pipeline, we considered some recommendations for analysis scripts in neuroimaging already present in the literature, grounded in software development (22). The design decisions made during development of our data processing modules offer a potentially valuable resource for other biologists in designing their own workflows. It is important to note that guidelines presented here do not always follow the sequential order of wet lab experiments; instead, they are presented in a non-chronological manner to better explain individual pipeline operations.

We provide a link to the GitHub repository housing the data processing pipeline^1^. As in daily work we use GitLab to host our code, the GitHub repository is a replica of it and contains sample (mock-up) data and metadata necessary to run the pipeline steps implemented in Python. Building upon the work of van Vliet and colleagues (22), we use examples from our workflow to illustrate each design principle, linking it with the scripts in the code repository (refer to the sections “Examples from the workflow”).

### 3.1 Choose the right technologies

The choice of technologies for pipeline implementation is largely dependent on the skills of team members. Teams are often diverse when it comes to expertise and preferred way of working; building upon this diversity can significantly enhance overall performance and problem-solving capabilities (23,24). While computational skills are becoming more prevalent among biologists (16), not all researchers are proficient in programming. Software with a graphical user interface (GUI) may help implement complex operations by eliminating the need to learn programming and has a shorter adoption period than code-based solutions (25).

**Examples from the workflow**

Our modules are implemented in Knime and Python, but designed in a manner to ensure interoperability (refer to section 3.4 for further explanation). Knime is among one of the most user-friendly workflow management systems and offers a wide array of learning resources (4); Python is currently considered the most popular programming language (19), well-suited for biology applications (20) and easy to learn. However, other technologies can be used with similar success when designing custom pipelines. The skills and preferences of team members should guide the selection of an appropriate technology solution.

### 3.2 Translate Tasks to Modules

Workflow tasks, once defined, are then converted to modules. Modules are the building blocks of a data processing pipeline, whether they be individual Python scripts or Knime workflows. By design, a module should perform a simple, singular task. To prevent scripts from becoming difficult to understand, complex tasks should be divided into smaller tasks across multiple modules. The general principle is to have each module be self-contained and run with minimum dependence on other modules. This way, any changes or issues can be addressed without disrupting other modules in the pipeline.

Our pipeline follows the logic of the wet lab experiments – it is organized in modules that correspond to experimental flow on the bench (see Figure 2). Since the workflow is hierarchical (i.e., each wet lab step relies on the results of the previous step(s)), we alternate wet lab experiments with data processing, rather than running the entire pipeline when all experiments are completed. Thus, the modularity of each data processing step is defined by related wet lab experiments. For example, a module can: i) analyze experimental readouts, ii) prepare pipetting schemes for the next experiment, iii) perform sample selection based on predefined criteria, iv) assemble machine-readable files for automated wet lab experiments, and v) generate a file for data storage in the database. Our experience shows that, for hierarchical pipelines, following the wet lab criterion is the optimal way to organize modules.

**Figure 2.**
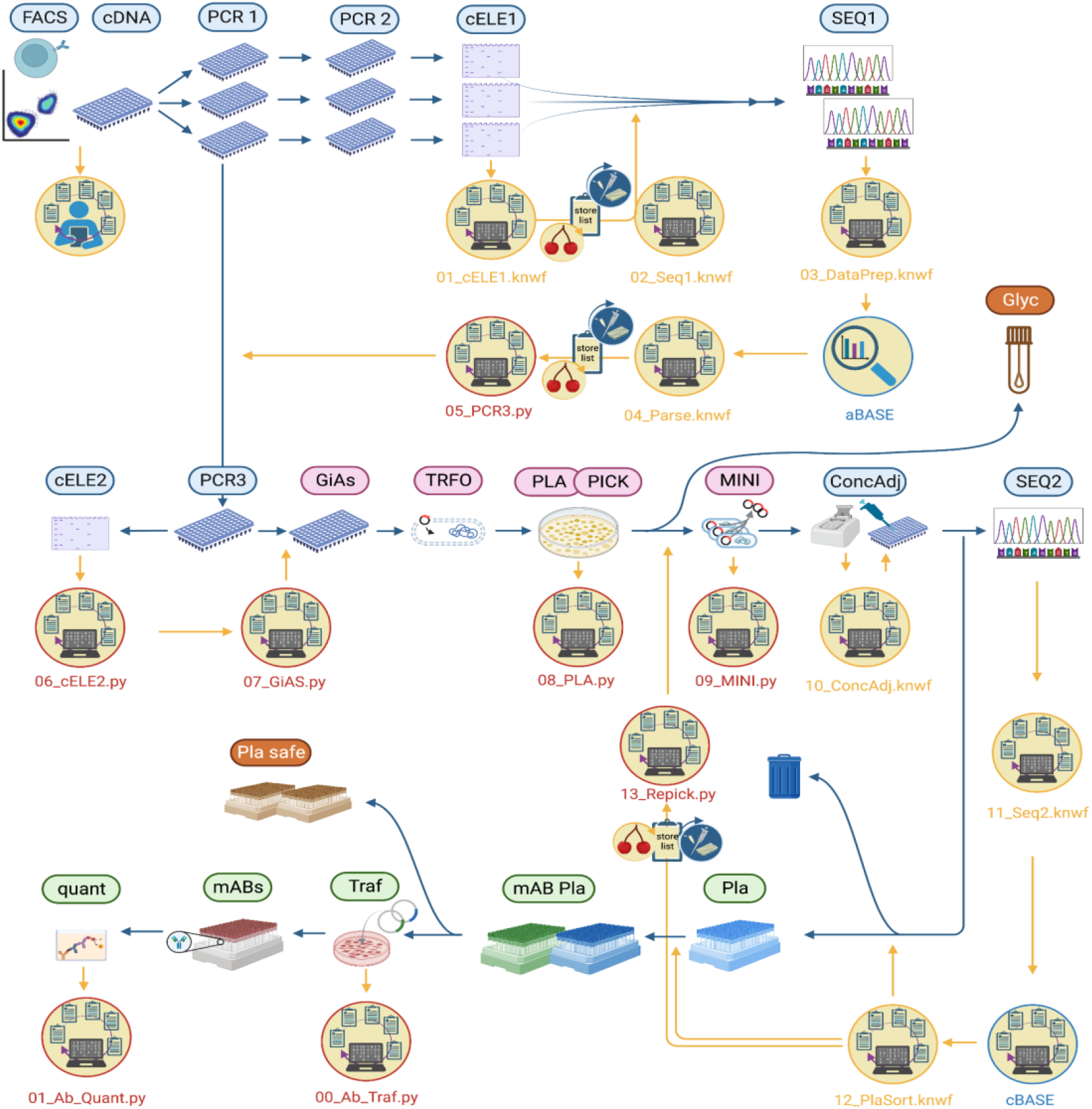
The workflow for the production of mABs from patient-derived individual B-cells: wet lab work steps (<u>icons with header</u>) and data processing steps (<u>yellow icons</u>). Sample flow is depicted by blue lines and information flow is depicted by yellow lines. The experimental procedure starts with FACS sorting of individual B-cells. cDNA is generated from individual B-cells, and PCR1 and PCR2 serve to amplify antibody chains. Successfully amplified chains, evaluated by capillary electrophoresis (<u>cELE1</u>), are sequenced (<u>SEQ1</u>). Upon positive sequence evaluation with the aBASE software (19), specific primers are used for the final amplification of the specificity-determining chain parts (<u>PCR3</u>). Amplification is quality controlled by electrophoresis (<u>cELE2</u>). The specificity-determining regions are cloned by Gibson Assembly (<u>GiAs</u>) into plasmids encoding the constant parts of the chains. Plasmids are amplified by transformed (<u>TRFO</u>) bacteria after plating (<u>PLA</u>) and picking (<u>PICK</u>) individual bacteria clones. Prior to quality control by sequencing (<u>SEQ2</u>), amplified plasmids are isolated (<u>MINI</u>), and plasmid concentration is measured and adjusted (<u>ConcAdj</u>). Based on sequencing results, functional plasmids (<u>PLA</u>) are sorted. Matching plasmid pairs (<u>mAB Pla</u>) are prepared for transfection (<u>Traf</u>) into HEK cells. mABs produced and secreted by HEK cells are harvested (<u>mABs</u>), and mAB concentration is quantified (<u>quant</u>). Glycerol stocks (<u>Glyc</u>) of transformed bacteria and plasmid aliquots (<u>Pla safe</u>) serve as safe stocks of plasmids. The colors of the wet lab work step captions indicate different workflow sections: blue – molecular biology; red – microbiology; green – cell biology. The colors of the data processing circle contours and font indicate the utilized technology: red – Python; yellow – Knime; blue – third-party software.

**Examples from the workflow**

To determine whether samples were successfully amplified in the final PCR step (PCR3, see Figure 2), we perform capillary electrophoresis (cELE2). The results of cELE2 are parsed by the module 06_cELE2.py. The module [loads experimental readouts](https://github.com/Malwoiniak/mABpy/blob/7d211be90fa05d94e27a2fe646abcca414ddc2db/Scripts/06_cELE2.py#L37) and [selects a set of samples](https://github.com/Malwoiniak/mABpy/blob/7d211be90fa05d94e27a2fe646abcca414ddc2db/Scripts/06_cELE2.py#L48) for further processing based on a band size threshold (in base pairs). The next module, 07_GiAS.py, takes up successfully amplified samples to prepare a [pipetting protocol](https://github.com/Malwoiniak/mABpy/blob/7d211be90fa05d94e27a2fe646abcca414ddc2db/Scripts/07_GiAS.py#L105-L124) for cloning by Gibson Assembly (GiAs) and a [database import file](https://github.com/Malwoiniak/mABpy/blob/7d211be90fa05d94e27a2fe646abcca414ddc2db/Scripts/07_GiAS.py#L95) (see Figure 2). For reference, we provide examples of raw [experimental readout](https://github.com/Malwoiniak/mABpy/blob/7d211be90fa05d94e27a2fe646abcca414ddc2db/Experimental_Data/cELE/baoscelep00380/2023%2002%2007%2014H%2041M%20Peak%20Table.csv) after capillary electrophoresis, [pipetting protocols](https://github.com/Malwoiniak/mABpy/blob/7d211be90fa05d94e27a2fe646abcca414ddc2db/Scripts/07_GiAS.py#L105-L124) (for automated liquid handling) and a [database import file](https://github.com/Malwoiniak/mABpy/tree/7d211be90fa05d94e27a2fe646abcca414ddc2db/07_GiAS_Out/07_GiAS_imports). Although our workflow is semi-automated and we often deal with machine-readable files, the modules are adaptable to manual workflows.

For example, the number of samples to be handled in the next workflow step can be easily modified by [defining the size of chunks](https://github.com/Malwoiniak/mABpy/blob/7d211be90fa05d94e27a2fe646abcca414ddc2db/Scripts/07_GiAS.py#L100-L102) in which samples are processed further. Since this workflow step operates on multiple 96-well plates, we set the number of samples to the batches of 96, but a lower number can be chosen for manual, low-throughput workflows.

### 3.3 Allocate separate computational space to modules

Each module should run in a separate computational space; that is, the input and output files generated by modules should be saved in dedicated directories. Ideally, script files (.py) reside in a separate directory, too. This computational environment needs to be accessible to all team members, must be backed up for data security, and should allow for seamless integration with different modules. These requirements are fulfilled by simple Network Attached Storage (NAS) drives, institutional set-up servers or cloud services. Within allocated computational space, the folder structure ideally reflects the architecture of the data processing pipeline.

**Examples from the workflow**

We created a dedicated computational workspace with a folder structure that resembles data processing steps. Keeping modules physically separated in dedicated directories allows i) deciding on and adhering to rigorous organization of both wet lab and data processing steps, resulting in better discipline; ii) easily accessing the resources (input files) required by each module without interfering with other modules; iii) demarcating Python from Knime modules. The published GitHub repository preserves the folder structure of Python modules in our pipeline. For example, directories 05_PCR3_Out to 09_MINI_Out are dedicated to wet lab steps from specific PCR (PCR3) to plasmid isolation (MINI), while the directory 01_Ab_Quant_Out is reserved for quantification of produced mABs (wet lab steps mABs and quant, see Figure 2). There is a separate folder for modules [manipulating bacterial colony information for repicking](https://github.com/Malwoiniak/mABpy/tree/7d211be90fa05d94e27a2fe646abcca414ddc2db/13_Repicks_Out) or for [input files used by modules](https://github.com/Malwoiniak/mABpy/tree/7d211be90fa05d94e27a2fe646abcca414ddc2db/Input). For reference, we also provide an [image representation](https://github.com/Malwoiniak/mABpy/blob/89620b9fcdb5949640ed8579a7527359b7db95a7/img/dir_str.png) of an in-depth listing of directories and files for Python modules.

We implemented a backup rotation scheme that involves daily (with a retention period of one week), monthly and yearly backups on external server. To safeguard against data loss in case of the server failure, the database is also backed up on a dedicated database server twice a day.

### 3.4 Define input and output files

Interoperability between modules (i.e., the ability of modules to work together seamlessly) should be given high priority when designing a pipeline. This ensures that data is passed between modules without loss of information or format, and each module can be developed and maintained independently while still functioning cohesively within the larger workflow system.

To achieve this, we defined input and output files that exchange data between modules. We refer to these as *intermediate files*, as they are created as part of a larger process and are not a final output of the workflow. Intermediate files provide standardized structure and syntax for exchanged data; they also serve as interfaces, i.e., files generated as output by one module can be then used as input by subsequent modules.

Storing intermediate files has several advantages: i) when errors occur, it is possible to rerun only the modules that failed, instead of running the entire pipeline again; ii) manual inspection of data is possible, which helps track the progress of the workflow and troubleshoot issues as they arise; iii) the autonomy of each module is maintained, which decreases the complexity of the pipeline and guarantees that modules run independently without relying on data saved in memory by other modules (22).

**Examples from the workflow**

Modules in our pipeline rely on one or more intermediate files, which are generated and saved by preceding modules. Intermediate files are often used by modules jointly with database export files, pipetting schemes or experimental readouts. For example, the module 08_PLA.py, which links the information on cloning by Gibson Assembly (<u>GiAs</u>, see Figure 2) with transformation and plating of bacteria samples (<u>TRFO</u> and <u>PLA</u>), starts by [loading](https://github.com/Malwoiniak/mABpy/blob/89620b9fcdb5949640ed8579a7527359b7db95a7/Scripts/08_PLA.py#L46-L49) the [pipetting schemes](https://github.com/Malwoiniak/mABpy/tree/89620b9fcdb5949640ed8579a7527359b7db95a7/Experimental_Data/qpix) (carried out by an automated colony picker) and [database export](https://github.com/Malwoiniak/mABpy/blob/89620b9fcdb5949640ed8579a7527359b7db95a7/Input/Gibson_IDs.xlsx) file to generate the [intermediate file](https://github.com/Malwoiniak/mABpy/blob/89620b9fcdb5949640ed8579a7527359b7db95a7/08_PLA_Out/Intermed_Files/PICK_Intermed_20230308-171741.xlsx). The latter is taken up by the next module, 09_MINI.py, which links bacteria with isolated plasmids. The module [loads](https://github.com/Malwoiniak/mABpy/blob/89620b9fcdb5949640ed8579a7527359b7db95a7/Scripts/09_MINI.py#L53-L55) the intermediate file and [creates](https://github.com/Malwoiniak/mABpy/blob/89620b9fcdb5949640ed8579a7527359b7db95a7/Scripts/09_MINI.py#L81) the [import](https://github.com/Malwoiniak/mABpy/blob/89620b9fcdb5949640ed8579a7527359b7db95a7/09_MINI_Out/MINI_import_20230309-165154.xlsx) for the database.

Although these steps are automated in our workflow, similar logic can be applied to manual workflows. For example, the module 08_PLA.py can take up any list of samples handled in low throughput. The number of samples to be processed is automatically deducted by the module from the list of samples, and the subsequent module would infer the number of samples from the intermediate file.

## 4. Dedicated databases allow solid data documentation and efficient sample tracing

The complexity of workflows determines the demands for its documentation and sample traceability. While simple workflows that handle some dozens of samples can likely be documented using spreadsheets, complex workflows that run multistep wet lab experiments for over a hundred samples require more advanced documentation, ideally in dedicated databases. A customized database facilitates retrieval of correct information between workflow end-points and saves significant time and effort otherwise spent on sample backtracking.

We chose FileMaker Pro Advanced (FM) as the database backend. FM is a low-code relational database management system that enables fast database creation and modification through drag-and-drop functionality (26). Since the database engine can be accessed through a GUI, querying the data is straightforward and queries are easily modifiable. Although FM may require some initial effort to become proficient, it remains an intuitive and user-friendly tool for biologists without prior experience in database design (27–29).

FM also provides scripting functionality which we used to automate the import and export of data for further processing in downstream applications. Customizable data exports facilitate quick quality control of the samples. This, in turn, is crucial for troubleshooting and for making decisions on the sample’s fate at key workflow steps.

**Examples from the workflow**

Our database is built around a concept of multiple, interconnected tables that store sets of unique records (here: information about sample state in the workflow). Records are connected through a series of relationships between individual tables; their uniqueness is guaranteed by a Universally Unique Identifier, a unique, 16-byte string assigned automatically to each record on data entry. Each subsequent, related table inherits the value of the record’s unique ID from a previous table, allowing the retrieval of sample information at any workflow step (see Figure S2). The data is stored in 19 distinct, interrelated tables.

Structuring the data model in multiple, interrelated tables minimizes the storage of redundant data. Information about a particular sample state is entered into the database only once, and then linked to other related data points as needed. This can help to reduce errors associated with redundant data entry, and save time and effort. Additionally, it ensures data consistency across multiple experiments, as the same sample information can be reused across different projects and analyses.

For example, we keep the information of bacteria plating separate from picking information. This allows picking of a new bacteria colony without the need to enter the same bacteria plate again to the database. Since bacteria plates are already linked to other information about the samples on previous wet lab steps (see Figure 2, steps from <u>FACS</u> to <u>TRFO</u>), connecting a repicked colony to plating information automatically handles the connection to other data in the backend (see Figure S3). Finally, our database provides relative flexibility, allowing the addition of new tables and seamless integration of experimental readouts. For example, the structure can be extended by adding the results of functional antibody assays to the information on harvested antibodies and linking a new table through shared IDs (see Figure S4).

## 5. Further guidance

It is important to consider design decisions that extend beyond building functional pipelines, and which can also enhance efficiency, organization and reproducibility of existing ones. These include using configuration files, optimizing sample handling by processing in batches, and more. Here, we discuss additional guidelines that can be applied to improve the performance and management of workflows beyond basic functionality.

### 5.1 Configuration files to reduce repetitive code

Rooted in software development, the DRY (“don’t repeat yourself”) principle implies minimizing code duplication (22). Parameters (e.g., file or directory paths, hardware-specific parameters) are often shared between modules, and one way to make them available is to create a configuration file that consolidates all defined parameters in a single location. Modules that require the parameters can import them from the configuration file by calling the unique variable name. There are several advantages to this approach – not only is it consistent with good programming practices, but it also reduces errors and saves time, as modifications to parameters need to be made only in the configuration file, rather than in multiple locations throughout the code (22). To minimize the possibility of variable duplication, we check whether a variable has already been defined in the configuration file before assigning it a value in a module. This way, we ensure that variables are used consistently, preventing unintended overwriting of data. Adhering to DRY principles makes the modules relatively resistant to modifications. All in all, using configuration files can streamline the process of adjusting parameters and settings across multiple modules.

**Examples from the workflow**

For Python modules, we provide the [configuration file](https://github.com/Malwoiniak/mABpy/blob/89620b9fcdb5949640ed8579a7527359b7db95a7/Scripts/config.py) (config.py file) that contains [file and directory paths](https://github.com/Malwoiniak/mABpy/blob/89620b9fcdb5949640ed8579a7527359b7db95a7/Scripts/config.py#L10), [experimental parameters](https://github.com/Malwoiniak/mABpy/blob/89620b9fcdb5949640ed8579a7527359b7db95a7/Scripts/config.py#L88-L92), [spreadsheet metadata](https://github.com/Malwoiniak/mABpy/blob/89620b9fcdb5949640ed8579a7527359b7db95a7/Scripts/config.py#L113-L115), and more. Each module imports only configuration parameters required for its run. For example, module 13_Repick.py imports the paths and metadata necessary for creation of a list of bacteria colonies to be re-selected for further processing.

To define file and directory paths, we employed the built-in *os* Python module. During the runtime, the paths are deducted from the module’s location. For example, module 00_Ab_Traf.py starts by [importing](https://github.com/Malwoiniak/mABpy/blob/89620b9fcdb5949640ed8579a7527359b7db95a7/Scripts/00_Ab_Traf.py#L11-L12) necessary paths from the config file and [determines](https://github.com/Malwoiniak/mABpy/blob/89620b9fcdb5949640ed8579a7527359b7db95a7/Scripts/00_Ab_Traf.py#L44) the location of directories and files [relative to the current (working) directory](https://github.com/Malwoiniak/mABpy/blob/89620b9fcdb5949640ed8579a7527359b7db95a7/Scripts/config.py#L12-L13) of the module. This makes the entire pipeline operating system-independent. Although we run the modules from a remote location, they can also be run locally if the folder’s structure remains unchanged. The hierarchy of the directories, as well as their names, can be adjusted to one’s needs by simply modifying the variables in the configuration file.

Similarly, any changes made to metadata of intermediate or database files can be updated in the configuration file, eliminating the need to modify multiple locations. As an example, modifying the metadata of the files required by an acoustic liquid dispenser to prepare samples for the final PCR run (<u>PCR3</u>) can be done easily by [updating the configuration file](https://github.com/Malwoiniak/mABpy/blob/89620b9fcdb5949640ed8579a7527359b7db95a7/Scripts/config.py#L102-L103), which is useful in case of changes in software.

A similar principle applies to Knime modules, where a separate configuration file specifies file and directory root paths for all the local machines that have access to Knime workflow. A custom Knime node reads the configuration file and adapts the path as necessary.

In adherence to the DRY principles, we have organized the utility functions – those that can be utilized by multiple modules for similar tasks – into Python files that contain only function definitions (i.e., no executable code). This approach allows for greater organization, reusability, and maintainability of code. The function files reside in [a designated folder](https://github.com/Malwoiniak/mABpy/tree/89620b9fcdb5949640ed8579a7527359b7db95a7/Scripts/utils) and are organized thematically. For example, the file microbiology_supporters.py contains the functions to assist with data processing on microbiology workflow steps, while the file plt_manipulators.py contains functions handling [manipulations of the plate grids](https://github.com/Malwoiniak/mABpy/blob/89620b9fcdb5949640ed8579a7527359b7db95a7/Scripts/utils/plt_manipulators.py#L54-L63) and [processing samples in batches](https://github.com/Malwoiniak/mABpy/blob/89620b9fcdb5949640ed8579a7527359b7db95a7/Scripts/utils/plt_manipulators.py#L99-L106), among others.

### 5.2 Efficient sample batching

Grouping samples together for batch processing is crucial for efficient use of resources, be it reagents, equipment or manpower. Sample batching can save the overall time and cost of the production process while reducing variability between samples and improving quality control and scalability of the workflows.

The production of mABs from individual B-cells is a multistep process, with quality assessment being performed after each wet lab step. Samples that do not meet the quality criteria are excluded from further processing, and only successful samples are selected for the next step of the workflow. Since batches of samples for each wet lab step are constantly updated, it is essential to keep track of batches already pushed forward in the workflow and of those still waiting for their turn.

In our workflow, wet lab experiments are conducted in batches of either 96 or 384 samples. To accommodate samples that cannot be processed due to limited space on the 96- or 384-plate grid, we implemented *store lists* to keep track of the leftover samples. Store list files are updated every time new samples are advanced in the workflow, with the priority given to *older* samples. This ensures that the ones that have been waiting the longest are processed first.

**Examples from the workflow**

An example of a store list for sample batching is created by the module 05_PCR3.py that prepares batches of samples for the final PCR step (PCR3). The module starts by [loading](https://github.com/Malwoiniak/mABpy/blob/89620b9fcdb5949640ed8579a7527359b7db95a7/Scripts/05_PCR3.py#L105-L106) the current [store list file](https://github.com/Malwoiniak/mABpy/blob/89620b9fcdb5949640ed8579a7527359b7db95a7/PCR3_Store_Lists/PCR3_StoreList_20230713-120500.xlsx) (updated previously by Knime module) and automatically [calculates the number of 96-sample batches](https://github.com/Malwoiniak/mABpy/blob/89620b9fcdb5949640ed8579a7527359b7db95a7/Scripts/05_PCR3.py#L115-L116) to be processed in PCR3 step. Then, the batches of selected samples are [pushed forward to the next workflow step](https://github.com/Malwoiniak/mABpy/blob/89620b9fcdb5949640ed8579a7527359b7db95a7/Scripts/05_PCR3.py#L145-L171), and the leftover samples are [saved](https://github.com/Malwoiniak/mABpy/blob/89620b9fcdb5949640ed8579a7527359b7db95a7/Scripts/05_PCR3.py#L118-L124) as a new store list file.

Computing the number of sample batches automatically is beneficial, as it minimizes user input, reducing the risk of human error. Additionally, we use *argparse*, a Python library for parsing command-line arguments, to get the plate barcodes for wet lab experiments (here: <u>PCR3</u>, <u>cELE</u> and <u>GiAs</u>, see Figure 2). The user inputs the latest plate numbers used on that workflow step as command-line arguments; the module [parses those as arguments](https://github.com/Malwoiniak/mABpy/blob/89620b9fcdb5949640ed8579a7527359b7db95a7/Scripts/05_PCR3.py#L83-L91) and [automatically assigns new barcodes](https://github.com/Malwoiniak/mABpy/blob/89620b9fcdb5949640ed8579a7527359b7db95a7/Scripts/05_PCR3.py#L126-L131) to current sample batches.

While our workflow is semi-automated and machine-dependent, similar principles can be applied to manual workflows. The number of samples to be processed in a single experiment can be [adjusted](https://github.com/Malwoiniak/mABpy/blob/89620b9fcdb5949640ed8579a7527359b7db95a7/Scripts/05_PCR3.py#L115) as needed by [modifying the size of a batch](https://github.com/Malwoiniak/mABpy/blob/89620b9fcdb5949640ed8579a7527359b7db95a7/Scripts/config.py#L84).

### 5.3 Getting feedback from modules

Because detecting errors that arise during module execution in complex workflows can be challenging, the modules should provide feedback whenever possible. Getting feedback from modules refers to collecting information from each step of the workflow to assess its performance and identify potential issues as they occur (22). Providing feedback enables collaborative usage of the pipeline, including by team members who may not be as familiar with all the individual operations performed by modules.

Feedback strategies can be satisfied by user interface features within the software. Most software offers visual cues and pop-up messages to inform users about their actions. For example, FM provides import log files and notifications on the import execution. Knime allows real-time assessment of computing operations through icons, indicating whether the run was successful. The result of each node’s operation can also be displayed upon request, enabling interactive troubleshooting. Another feedback strategy to consider is input validation; FM provides options of validating the IDs to be unique, non-empty and unmodifiable, and retrieves the messages on failed import if these conditions are not met. Similar validation can be implemented in Python, where incorrect user’s input can trigger clear and informative error messages that show what went wrong and suggest possible solutions. Additionally, a good approach to automated and recurring feedback is saving custom files that include run parameters or any other information helpful for potential troubleshooting or documentation. As a minimum requirement though, in case of scripted modules, simple printout messages help to orient the users about the status of the run.

**Examples from the workflow**

We have included printout messages for key steps in Python modules. For example, module 05_PCR3.py provides information on the total [number of samples (and 96-well plates) to process in final PCR step, number of leftover samples saved as a store list file, together with the directory and filename of the store list](https://github.com/Malwoiniak/mABpy/blob/89620b9fcdb5949640ed8579a7527359b7db95a7/Scripts/05_PCR3.py#L116-L124). Further, the *argparse* module is employed in most Python scripts. It automatically generates help, usage and error messages, which prevents incorrect input from being passed during the run.

For Knime modules, we implemented self-documenting features, where a custom file is generated upon the module run, containing the metadata such as run parameters, generated input and output files, user data, timestamps or messages. Together with the intermediate files saved at workflow steps, self-documentation features provide frequent feedback on the progress of data processing steps.

### 5.4 Organizing the pipeline

To maintain the organization of the pipeline, we recommend introducing a system that is user-friendly and not too complex, so that the users are encouraged to adhere to continuous documentation. It is a good practice to separate modules that are part of the main computational pipeline from auxiliary ones or those that represent work in progress. Regular inspections and a simple cleaning strategy for modules can ensure workflow maintenance with minimal effort (21).

A manual should be provided for each module in the pipeline to ensure proper usage and minimize the likelihood of errors due to incorrect execution. Ideally, all factors that could affect the pipeline execution should be documented. This allows for replication of the analysis in the future by yourself or others. Each module should have a designated champion who understands the module’s architecture and logic and can assist with troubleshooting and implementing the changes. In addition to a well-defined folder structure for hosting the modules, we recommend implementing a Version Control System (VCS) to track modifications to modules that are part of the main pipeline. Finally, adhering to a comprehensive documentation scheme aids in the maintenance of the workflow.

Writing detailed manuals, documenting module logic, and implementing version control require discipline, but contributes to efficient maintenance of complex pipelines. Even routine operations can require consulting the documentation; in our experience, the effort of creating manuals and adhering to version control pays off in sustaining the pipeline.

**Examples from the workflow**

Distinguishing between the main and auxiliary Python modules is achieved by keeping a coherent naming convention and shared directory. Python modules hold names starting with [a sequential, two-digit number](https://github.com/Malwoiniak/mABpy/tree/89620b9fcdb5949640ed8579a7527359b7db95a7/Scripts). The outputs of module runs are kept in separate directories, suffixed with _Out. Git is used as VCS to track changes to main Python modules, allowing for restoration of the previous module’s versions if needed. Manuals for each workflow step [are also tracked by VCS](https://github.com/Malwoiniak/mABpy/blob/92dc82fe5b5b4cd41f1d38e23348859e884e25b6/manuals/mABpy_tutorial.pdf).

We keep a documentation strategy for wet lab workflow and database imports by using Electronic Lab Notebook (ELN) software. Wet lab experiments, data imports, and modifications of database structure are recorded as separate entries in the ELN, which serves as a reliable record-keeping tool. Finally, we emphasize comprehensive code documentation, recognizing that code is read much more often than it is written (24). This approach helps us create well-documented modules.

## 6. Proof of Concept – Tailored Data Processing Pipeline and Database for Automated Stem Cell Culture (ASCC)

To showcase the versatility of our design, we applied the design principles to develop a data processing pipeline and a database for automated stem cell culture. Recently, we have established an automated cell culture platform that integrates a robotic liquid handling workstation for cultivation and differentiation of human induced pluripotent stem cells (hiPSCs). The hiPSCs are expanded and differentiated into brain microvascular endothelial cells (BMECs) for generation of an *in vitro* blood-brain barrier (BBB) model (30). Mature BMECs are seeded on TransWell plates for a 2D permeability BBB model. Trans-endothelial electrical resistance (TEER) is then measured to assess the integrity of the barrier (for a detailed protocol, refer to the work published by Fengler et al. (30)). BBB models generated in high-throughput scale with close-to-physiological characteristics can facilitate the screening of BBB-penetrating drugs, aiding the development of targeted drug delivery systems for neurological disorders (31).

Here, we present a proof-of-concept pipeline focused on hiPSCs differentiation into BMECs that is flexible and can be expanded to implement generation of other hiPSCs-derived cell types, including astrocytes, neurons, microglia or monocytes. The pipeline consists of Python scripts, FM database and FM scripts; below, we discuss the design process and how it aligns with the design principles.

### 6.1 Getting started – ASCC

We started the design by outlining:

1. The wet lab experiments (cycles of thawing, seeding and harvest of cells, see Figure S5).
2. The methods to document metadata, such as: i) hiPSCs batch information provided by a supplier; ii) culture conditions (including medium batch information and supplements); iii) the experiments’ timestamps; iv) quality control data (such as cell viability assays and cell counts); v) equipment settings (parameters controlled by automation); and vi) user annotations.
3. The repetitions or pausing steps in the workflow, i.e., interruptions of the differentiation process, including freezing of cells on differentiation stages for future experiments or on expansion stages to allow differentiation into other cell types.
4. The methods of documenting the workflow for human supervision and potential error detection.

### 6.2 Translate tasks to modules – ASCC

We then translated tasks to modules by following the wet lab criteria, making sure that each module performs a singular task and has little dependence on other modules (other than logical dependence that results from hierarchical nature of the workflow). Modules execute the following tasks: i) analyzing the experimental readouts (machine- or manually-generated), ii) parsing metadata, and iii) generating an import file for database storage.

During expansion and differentiation, iPSCs undergo seeding and harvest cycles across various plate formats (for example, 4-, 6- or 12-well plate grid for TransWell plates). Cell count and viability assays are performed on each harvest cycle or whenever an assessment of cell confluency is required. Module 00_cellcount_parser.py reads the plate format from user’s input and processes the file with cell count and viability information (generated by the automated cell viability analyzer, ViCell). During the entire process of cultivation, cells are imaged daily for quality control. Module 01_img_parser.py parses [the images](https://github.com/Malwoiniak/StemCellFactory/blob/319d97e2440487fe775c3bea5a64beeae23ce631/Scripts/01_img_parser.py#L56-L64) generated by a spinning disk confocal microscope and [creates a .json file with metadata](https://github.com/Malwoiniak/StemCellFactory/blob/319d97e2440487fe775c3bea5a64beeae23ce631/Scripts/01_img_parser.py#L65-L71) of all images linked to a given plate. On final WF steps, cells are seeded on TransWell plates for TEER measurements. Module 02_teer_parser.py parses the experimental readouts and [creates an import file for database storage](https://github.com/Malwoiniak/StemCellFactory/blob/319d97e2440487fe775c3bea5a64beeae23ce631/Scripts/02_teer_parser.py#L66-L69).

We also implemented the config.py file to store [file and directory paths, spreadsheet metadata](https://github.com/Malwoiniak/StemCellFactory/blob/319d97e2440487fe775c3bea5a64beeae23ce631/Scripts/config.py#L10-L46) and [configuration parameters](https://github.com/Malwoiniak/StemCellFactory/blob/319d97e2440487fe775c3bea5a64beeae23ce631/Scripts/config.py#L58-L92). [The utility functions](https://github.com/Malwoiniak/StemCellFactory/tree/319d97e2440487fe775c3bea5a64beeae23ce631/Scripts/utils) are grouped thematically: cellcount_img_tools.py handles .txt files generated by ViCell and [input_readers.py](https://github.com/Malwoiniak/StemCellFactory/blob/319d97e2440487fe775c3bea5a64beeae23ce631/Scripts/01_img_parser.py) parses the user input.

### 6.3 Allocate separate computational space to modules – ASCC

We created a designated computational space for modules, input files (including metadata, experimental readouts and files exported from the database) and output files, with an automated backup rotation scheme of daily, monthly and yearly backups. There are designated directories for database export files for [cell count](https://github.com/Malwoiniak/AutomatedStemCellCulture/tree/668dfe5766f0cc62d0344a15c41a89a486d20108/Input/plt_exports) and [image parsing](https://github.com/Malwoiniak/AutomatedStemCellCulture/tree/668dfe5766f0cc62d0344a15c41a89a486d20108/Input/medium_exports). [Confocal microscopy images](https://github.com/Malwoiniak/AutomatedStemCellCulture/tree/668dfe5766f0cc62d0344a15c41a89a486d20108/Input/img) (per plate) and [files generated by the automated cell viability analyzer](https://github.com/Malwoiniak/AutomatedStemCellCulture/tree/668dfe5766f0cc62d0344a15c41a89a486d20108/Input/experimental_data) are kept in individual folders. [Scripts, .json files](https://github.com/Malwoiniak/AutomatedStemCellCulture/tree/668dfe5766f0cc62d0344a15c41a89a486d20108/JSON) and [output files generated at each workflow step](https://github.com/Malwoiniak/AutomatedStemCellCulture/tree/668dfe5766f0cc62d0344a15c41a89a486d20108/4well_Out) also have separate directories.

### 6.4 Define input and output files – ASCC

We defined input and output files to share data between modules. Depending on the plate format and whether cells are in expansion or differentiation stage, the modules access different input files. For example, in an early expansion stage, the module 00_cellcount_parser.py takes up the [database export file](https://github.com/Malwoiniak/AutomatedStemCellCulture/blob/668dfe5766f0cc62d0344a15c41a89a486d20108/Input/plt_exports/4Well_export.xlsx) (automatically generated by a FM script) and [experimental readout file](https://github.com/Malwoiniak/AutomatedStemCellCulture/blob/668dfe5766f0cc62d0344a15c41a89a486d20108/Input/experimental_data/4well/Barcode4Well_1.txt) to generate the [import file](https://github.com/Malwoiniak/AutomatedStemCellCulture/blob/e6a42f3596b360d95d231dc1c99a7343a5c15e73/4well_Out/cell_count/20231107-155647_4well_import.xlsx) for database storage (saved with a timestamp). To facilitate the automatic imports to the database, records processed at the time are saved as [separate files](https://github.com/Malwoiniak/AutomatedStemCellCulture/blob/e6a42f3596b360d95d231dc1c99a7343a5c15e73/4well_Out/cell_count/IMPORT_NOW.xlsx), updated after each script run. Modules 01_img_parser.py and 02_teer_parser.py follow similar logic when processing confocal microscopy images and TEER measurement files, respectively.

### 6.5 Dedicated Database – ASCC

While conceptualizing ASCC database structure, we considered several factors:

1. The information should not be redundant, i.e., it should only be entered in the database once. If the same information is entered more than once, there is probably a need to restructure the database and keep the redundant information in a separate table. For example, [the information on the culture medium batch](https://github.com/Malwoiniak/AutomatedStemCellCulture/tree/1ea14c4066ad56432917e00d21a6983f4c4d86de/Medium) is registered only once and stored in a distinct table. Upon a medium change event, this information is populated by fetching the ID of the medium batch and adding it to the table storing the plate information (see Figure S6 (A)).
2. The tables are organized following the thawing, seeding, harvesting or freezing events, as they imply the initiation of a new process: e.g., change of the barcode or plate format, storing frozen cells in tanks, or collecting metadata, such as cell viability assays and cell count. For example, the information on harvesting differentiated BMECs from 6-well plates by pulling and seeding on multiple 12-well TransWell plates is kept in separate tables linked through a one-to-many relationship. This ensures that: i) any manipulations performed with the 6-well plate (medium change or cell count) are independent from those performed with the 12-well TransWell plates (TEER measurements or a well treatment); and ii) no redundant information is entered in the database, i.e., only the IDs of the 6-well plates are populated in the TransWell plate table (see the Figure S6 (B)).
3. We implemented separate tables also due to the different hiPSCs differentiation methods, each requiring distinct processes. Although the current ASCC database assumes BMECs differentiation, other cell types (e.g., astrocytes) will be considered in the future. In the current structure, new tables for astrocytes differentiation can easily be linked to existing ones (see the Figure S7).

## Discussion

The ability to document, process and analyze large datasets to identify new patterns and relationships has become increasingly important in modern biology. In the evolving landscape of biology research, it is not only large specialized laboratories that are embracing HT/HC techniques, but smaller biology labs are also progressively implementing similar methods. However, dealing with data generated in HT/HC schemes poses challenges related to sample backtracking, quality control, record keeping, and data curation and storage, among others. This can lead to compromised data integrity, potential data loss, and ineffective workflows. These challenges can be addressed by implementing modular data processing pipelines, which help to make substantial progress toward optimizing data management practices.

We employed the computational pipeline for data processing in a complex, semi-automated, multistep workflow for the production of mABs to tackle similar challenges and monitor all workflow steps, ultimately aiming to enhance data governance in our experimental setup. The design principles presented here can serve as guidance for the development of data processing pipelines in biology. The versatility of our approach allows for its application to diverse biological problems concerned with intensive data collection activities in a variety of settings, in which data undergoes continuous processing, analysis and modification before reaching the endpoint.

We showed how our design can help minimize the reliance on error-prone and resource-intensive manual data handling, significantly reducing errors and optimizing both time and resource usage. The modular nature of the design allows for flexibility to handle samples in high- and low-throughput settings. While the modules are designed to function in isolation, we combine them into a custom pipeline that operates on the mAB production workflow.

Our approach also helps streamline data processing and documentation, leading to faster generation of insights (allowing more time for other tasks), improving data reusability and promoting seamless data exchange. Additionally, we showed how deploying the dedicated database facilitates the standardization of data processing procedures. By implementing the tailored database, we established consistent and structured approaches to storing and organizing data, allowing us to efficiently track the samples throughout the workflow. Overall, adhering to the design principles outlined in this work can assist in enhancing the accuracy and fostering the reproducibility of data analysis by combining standardization, documentation, scalability, and error detection, and is a step towards more robust frameworks for data processing in biology.

Although our modules are tailored for mABs production, we foresee that our approach can support other biologists in building their own small-scale data processing pipelines for individual use. This approach is different from large-scale pipelines developed by bioinformaticians to manage massive volumes of data (32). Given that our team primarily consists of biologists, we do not have ready access to specialized support from computer scientists in our daily work. Thus, we focused our design efforts on feasibility for biologists with limited programming experience by employing software with GUI, such as Knime and FM, besides Python.

With some modifications, individual modules presented here can also be adapted by other biologists and incorporated into their own workflows. To that end, we provide a GitHub repository^2^ containing the mock-up data and the simplified pipeline, together with detailed instructions on how to download, install, and run it. Furthermore, we demonstrate how the code can be customized to accommodate data processing routines based on individual requirements^3^. This can serve as a starting point for experimental researchers in constructing their pipelines by reusing and modifying the provided code.

To test the applicability of our design to other settings, we applied the design principles to automated hiPSCs culture, proving that it was possible to adapt the framework to stem cell-related research. In the long run, we expect that this approach will be beneficial for ASCC, as it can help to: i) standardize and automate relevant data storage; ii) facilitate data-centric conclusions to gain novel insights; iii) make informed decisions for potential troubleshooting; iv) use data to optimize processes by systematic analysis through artificial intelligence or “design of experiment” approaches. As pipelines for continuous data processing and analysis are now essential in domains such as multi-omics data integration (33,34), HT/HC screening (35), data-driven modeling (36), as well as long-term environmental monitoring (37), evolutionary biology (38–40) or plant phenotyping (41), among others, we believe that design recommendations proposed here can find their target audience and be a source of inspiration to other researchers in developing their own data processing modules.

In the context of data-intensive science, there is a growing demand from various stakeholders, such as the scientific community, funding agencies, publishers and industry, for data to meet the standards of being Findable, Accessible, Interoperable, and Reusable (FAIR) over the long-term. This also concerns research processes beyond data, including analytical workflows and data processing pipelines. Many projects have already adopted different elements of FAIR principles into their data (and non-data) repositories (42).

Our data processing pipelines align with FAIR principles in several ways. First, we satisfy the *findability* facet of FAIR by making the mAB^4^ and the ASCC^5^ pipelines publicly available via GitHub and assigning a persistent identifier. Clear documentation in the current paper and in the GitHub repository provide the necessary information to understand the design decisions, data and processes involved in running the pipelines. To satisfy the *accessibility* facet, we include contact information and apply an open access policy to the code, granting unrestricted access to both the mock-up data and pipeline modules. Standardized file formats and clear naming conventions throughout the pipelines aid *interoperability* by keeping a consistent and organized approach to data representation. Moreover, using standardized data formats enables compatibility and facilitates data exchange across pipelines. To account for the *reusability* aspect of FAIR, we design modular pipelines, emphasizing the granularity of the modules. Published documentation defines how modules work and how they interact with input and output data. Finally, adhering to the outlined principles for data processing may foster compliance with FAIR principles during publication. This approach ensures that both data and metadata are well-structured, minimizing the effort to achieve data accessibility and findability upon publication.

One limitation of the system is partial reliance on licensed software, such as FM and GitLab. Nevertheless, open-source alternatives can be employed instead. For example, LibreOffice Base is an open-source, no-code database management software and offers similar functionality to that of FM, allowing users to create and manage databases using a graphical interface with features for creating tables, queries, and reports (43), making it a suitable option for researchers with little to no programming experience. Alternatively, if team members have some level of SQL knowledge, they can consider alternatives like MySQL for building and managing databases. MySQL Workbench offers a user-friendly interface to simplify database creation (44), and there are numerous resources available online to learn the fundamentals of SQL and get started.

Another interesting alternative to transform existing, code-based databases into interactive applications is NocoDB – an open-source, no-code platform that turns databases into spreadsheets with intuitive interfaces, making it possible for teams to create no-code applications. It supports MySQL, SQLite or PostgreSQL databases, among others, but it also provides the functionality of building databases from scratch (45). NocoDB is becoming popular within the community as an alternative to non-open-source solutions. As the platform gained attention from users seeking efficient and customizable data management solutions, there are plenty of online learning materials available, including tutorials, documentation and community forums.

Although we use the GitLab for Enterprise as a VCS to host the pipeline, the free GitLab version offers essential features for individual users and provides enough functionality to implement data processing pipelines (despite some storage and transfer limits). Alternatively, GitHub can be used as a VCS, as it has traditionally been more widely recognized and utilized in the developer community due to its extensive user base and integration options (46).

Finally, labs can also consider existing Laboratory Information Management Systems (LIMS) when automating the workflows to efficiently handle samples and their related data. LIMS are software applications used to streamline laboratory operations, sample tracking, data management and reporting, offering integration with various lab instruments (47). Although these systems provide several advantages over custom-designed pipelines, investing in a full-fledged, commercial LIMS can be expensive. LIMS are also lab-centric, that is, they are designed to cater to the overall needs of the laboratory as a whole and thus require tailoring to each individual lab. As the goal of LIMS is to create a centralized system that manages all laboratory activities and data, a significant effort has to be spent on specifying requirements for the LIMS systems.

Workflow-centric custom pipelines, on the other hand, are designed with a focus on optimizing and automating a particular laboratory workflow or a set of interconnected processes. Thus, our custom approach to modular pipeline development offers several benefits: it is cost-effective; can be tailored to specific needs; and allows rapid, iterative prototyping and experimenting with different solutions. What’s more, pipelines designed with this approach in mind are easier to maintain and can grow organically, adapting to changing requirements of individual projects. They can also be designed to integrate better with other custom systems already used by labs. Finally, with a custom pipeline the labs retain full control and ownership of the process, allowing for flexibility in response to changing needs and providing protective measures to the laboratory’s intellectual property.

## Supporting information

See the attached Table S1.xlsx file.

## Acknowledgments

We thank the Helmholtz Association for funding HIL-A03. All figures were created with BioRender.com

## Contributions

Conceptualization, methodology and software: M.K., D.S.; Data curation and investigation: S.F., M.K., B.K., A.M., D.S., M.S.; Resources: J.K., M.R., E.S., S.vH.; Validation: M.K., M.S., G.S.; Project administration and supervision: P.D., E.F., D.S.; Visualization: M.K. E.M. D.S.; Writing: J.B., E.F., M.K., D.S., G.S.; Funding Acquisition: P.D., E.F., J.K., L.K., H.P., D.S.

## Supplementary Material

**Figure S1.**
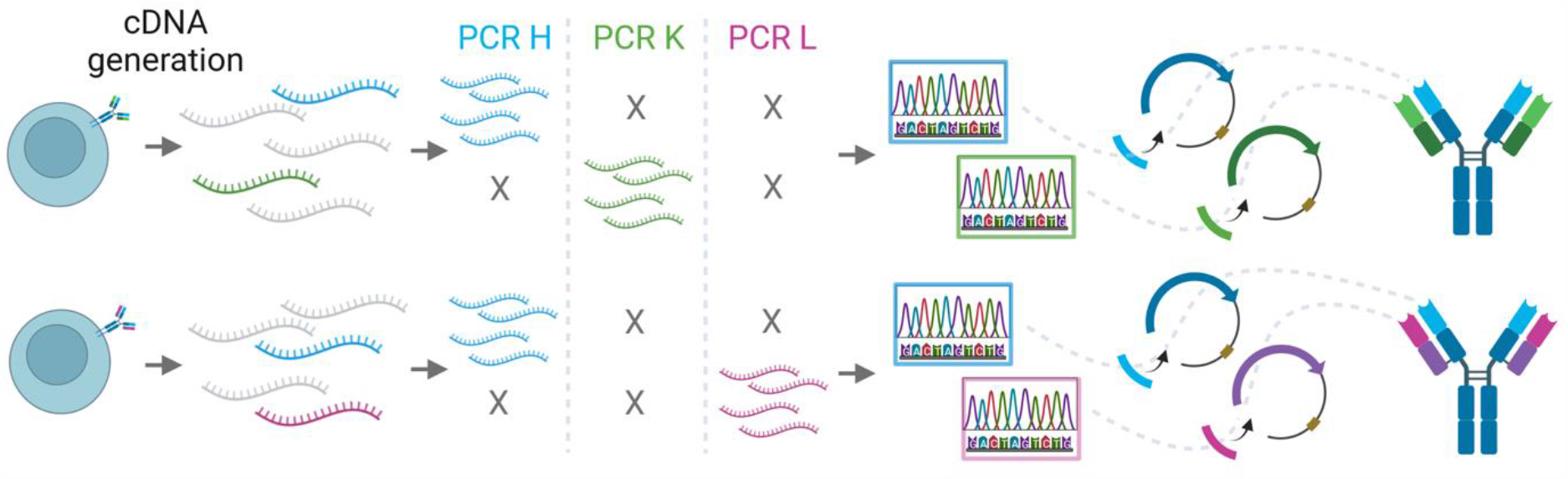
Cloning strategy for variable antibody regions. After cDNA generation, three parallel PCR reactions are performed to amplify the heavy chain variable region (light blue) and either the Kappa (light green) or Lambda (pink) variable region. Upon verification by sequencing, the variable regions are cloned into plasmids encoding the constant part of the respective antibody chain (dark blue: heavy constant part; dark green: Kappa light constant part; purple: Lambda light constant part). See Figure 2 for a comprehensive overview. For an overview on B-cell receptor and antibody variability refer to Figure 1 by Khatri et al (48) and Figure 1 by Mikocziova et al. (49).

**Figure S2.**
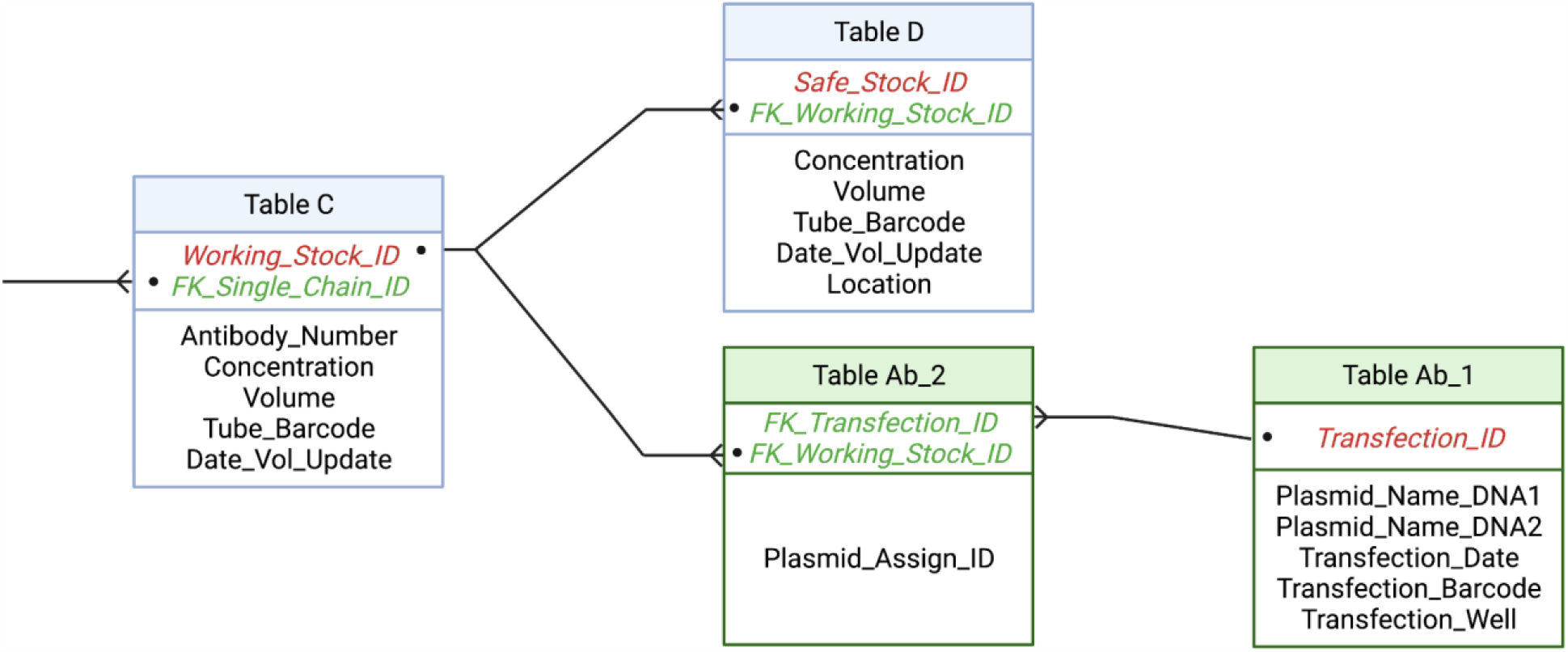
An excerpt from the database structure to explain its relational aspects. Table C stores information on isolated plasmids of paired antibody chains (heavy and light), and Table D is a repository of plasmid aliquots stored as safe stocks in separate (physical) locations for future reproduction of plasmids or downstream application. Table Ab_2 connects the information on plasmid pairs with subsequent transfection of HEK cells. Fields (i.e., columns) Working_Stock_ID, Safe_Stock_ID and Transfection_ID store unique IDs of each record per table. Fields starting with FK_ are records’ IDs inherited from related tables. Records (sample states) are connected through one-to-many relationships in the database structure; for example, a plasmid from Table C can be used for creation of a safe stock aliquot (in Table D) multiple times, while one aliquot of safe stock (in Table D) is associated with exactly one plasmid from Table C. Similarly, one transfected HEK cell sample (i.e., transfection well, in Table Ab_1) is associated with exactly two plasmids (a pair of heavy and light chain plasmids, in Table C), while this plasmid pair can be used for multiple transfections.

**Figure S3.**
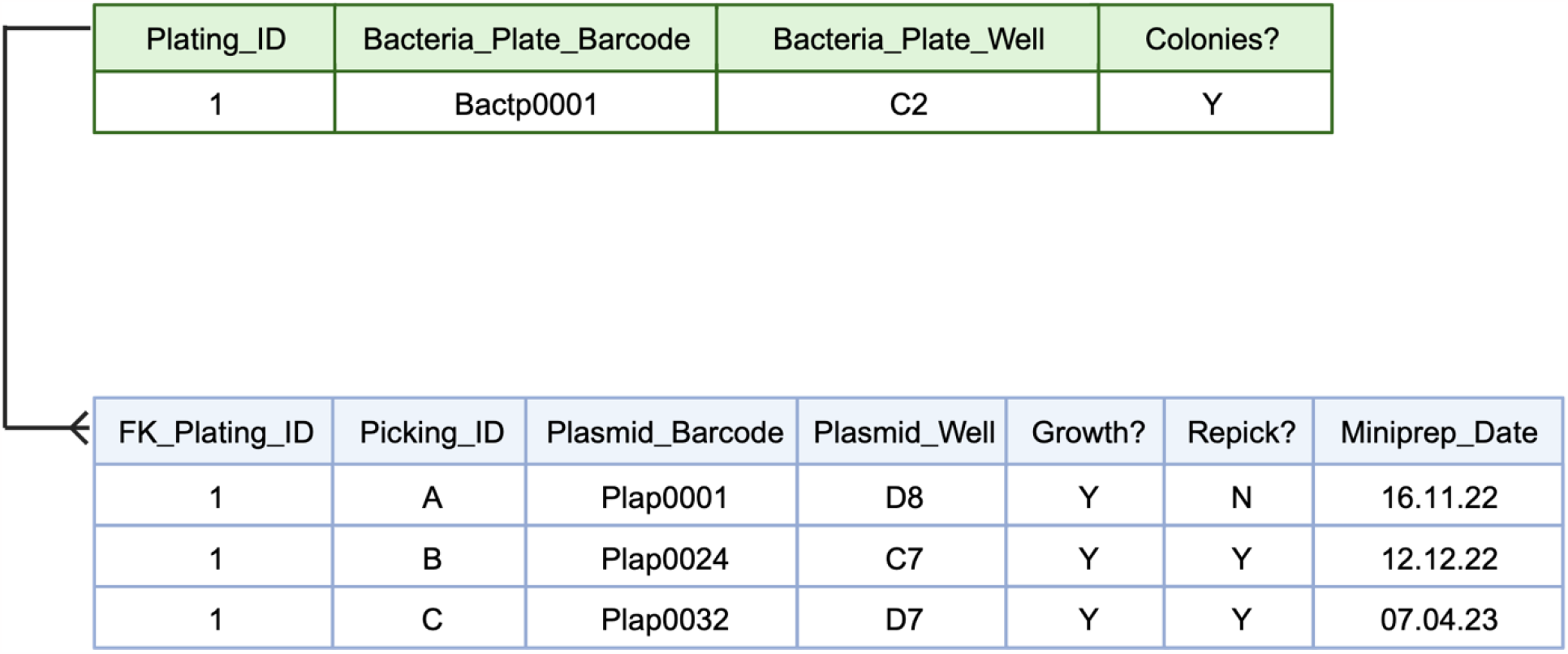
Simplified view of tables storing bacteria plate information (upper, green) and colony picking information (lower, blue). Picked colonies can be connected to already imported bacteria plate information through the identifiers (Plating_ID/FK_Plating_ID), reducing the redundancy of stored data. The identifier Plating_ID is unique in the bacteria plating table but not in the colony picking table, allowing for picking of multiple colonies from the same bacteria plate (one-to-many relationship in the database structure). The colony picking table has its own unique identifier (Picking_ID).

**Figure S4.**
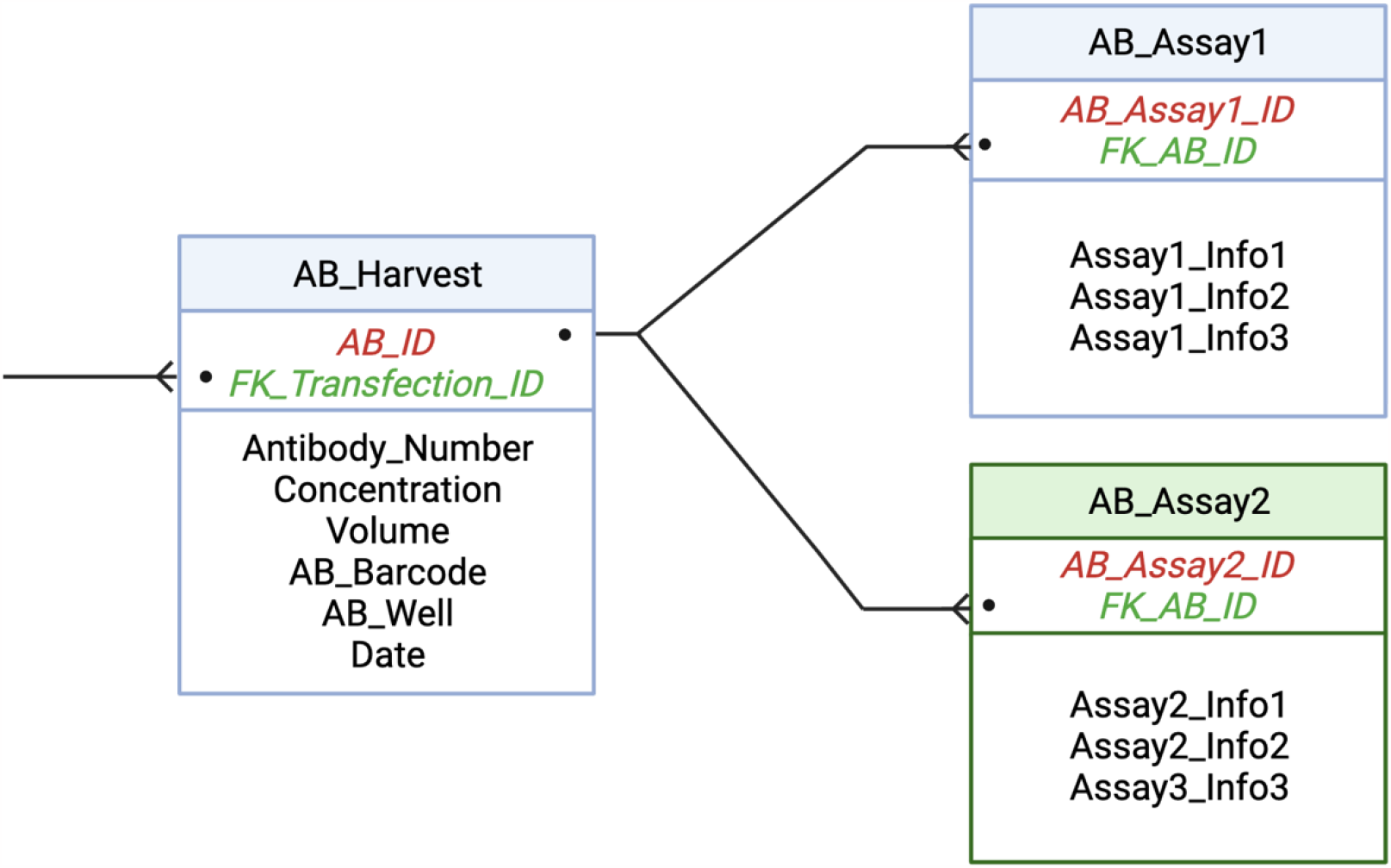
A hypothetical extension of database structure by table addition. The flexibility of the design enables smooth integration of new information (e.g., experimental readouts), which allows for efficient management of diverse datasets within the database. Here, the information on two hypothetical functional antibody assays (AB_Assay1 and AB_Assay2) were appended to the information on the harvested antibody through the AB_ID identifier, thus getting linked to any previous information on that sample (starting from the B-cell donor).

**Figure S5.**
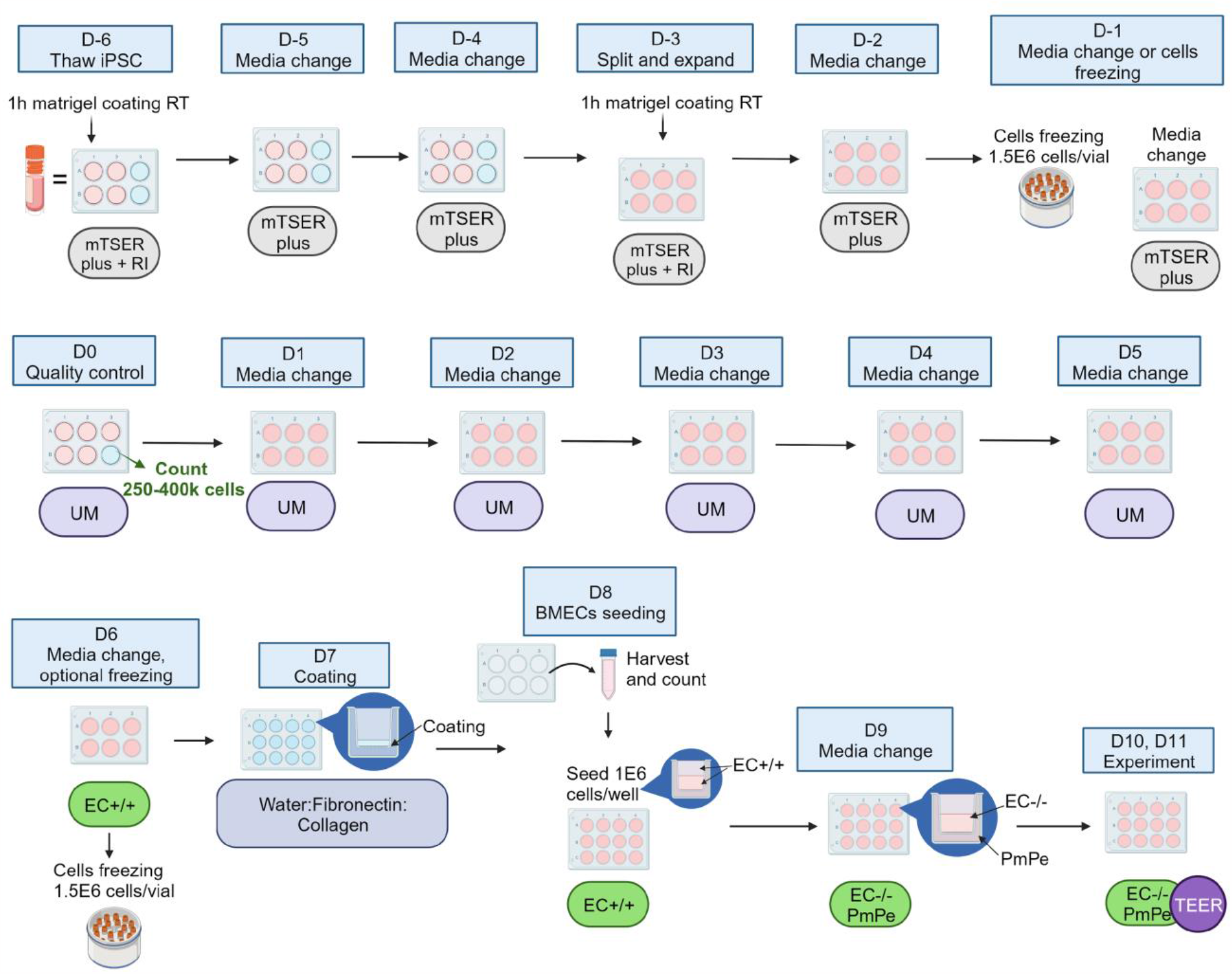
Wet lab steps of ASCC workflow: thawing, expansion and differentiation of iPSCs into BMECs in functional BBB model. Wet lab steps and timepoints (relative to the start of the differentiation – day 0: D0) are indicated by blue rectangles. Cycles of thawing (D-6), possible freezing (D-1, D6), harvest and seeding (D-3, D8) of cells. Count and viability assays are carried out on D-3, D0 and D8. Media used at each timepoint are indicated by ellipses – gray: mTSER plus medium with/without rock inhibitor; violet: Unconditioned Medium (UM); green: Endothelial Cell Medium with/without supplements (EC +/+, EC -/-, respectively). The TEER measurement timepoints (D10, D11) are indicated by a purple circle. For detailed protocol, refer to Fengler et al (30).

**Figure S6.**
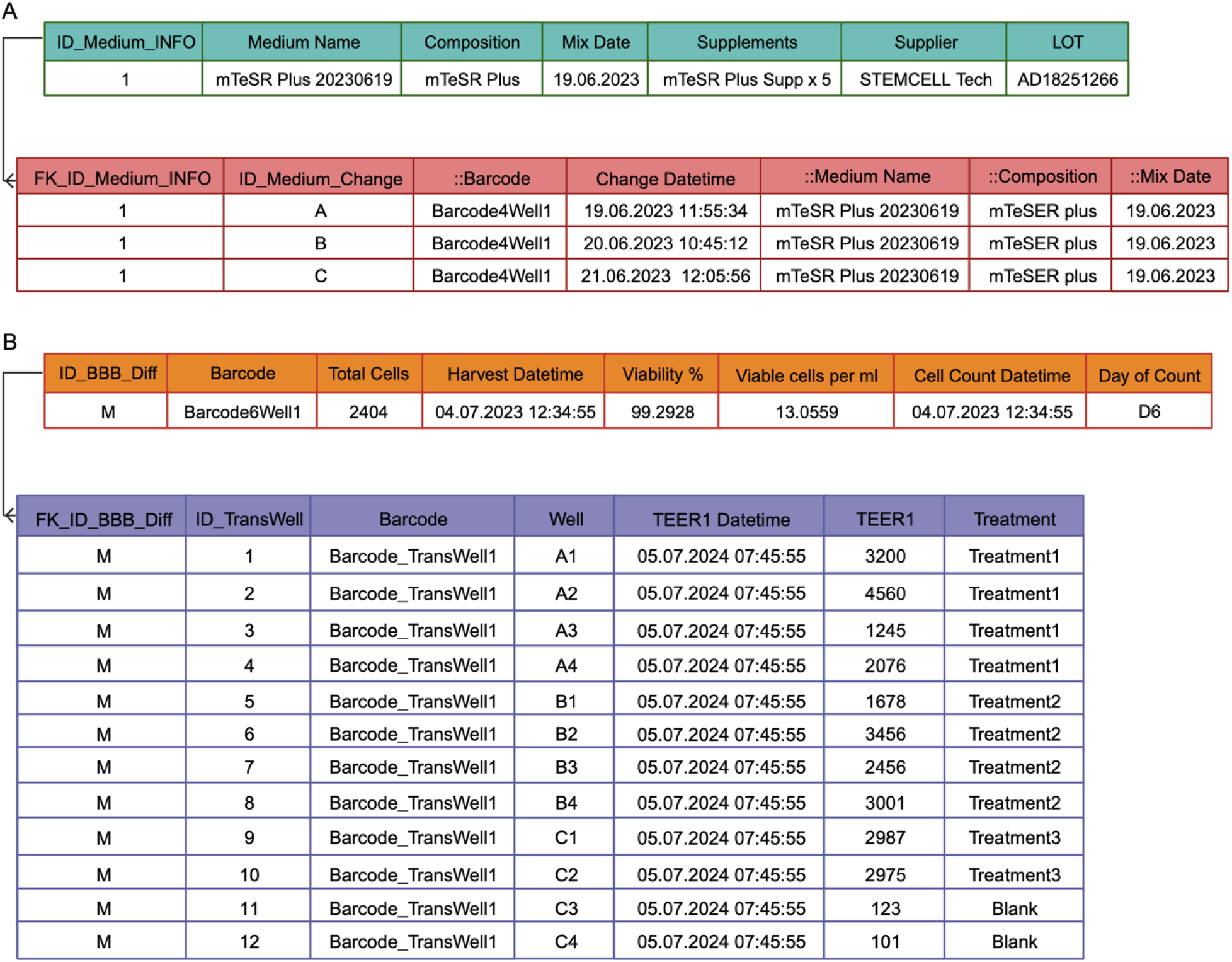
Conceptualizing ASCC database. **A)** No-redundancy principle. The information on each medium batch is stored in the database only once (upper table, green headers). Whenever a media change is performed, the unique identifier of the concerned medium batch (ID_medium_INFO) is fetched on the backend by a FM script and populated in the media change table (lower table, red headers). Information in columns ::Medium Name, ::Composition, and ::Mix Date is fetched from the medium table (upper table, green headers). Information in the column ::Barcode is fetched from another table (not shown). **B)** Example of the harvest event that guided the design of database structure: pulling differentiated cells from 6-well plate and seeding on 12-well TransWell plate. At this workflow stage, cells could also be frozen and stored for future experiments. Implementing a one-to-many relationship between 6-well and 12-well TransWell plates helps avoid storing redundant information.

**Figure S7.**
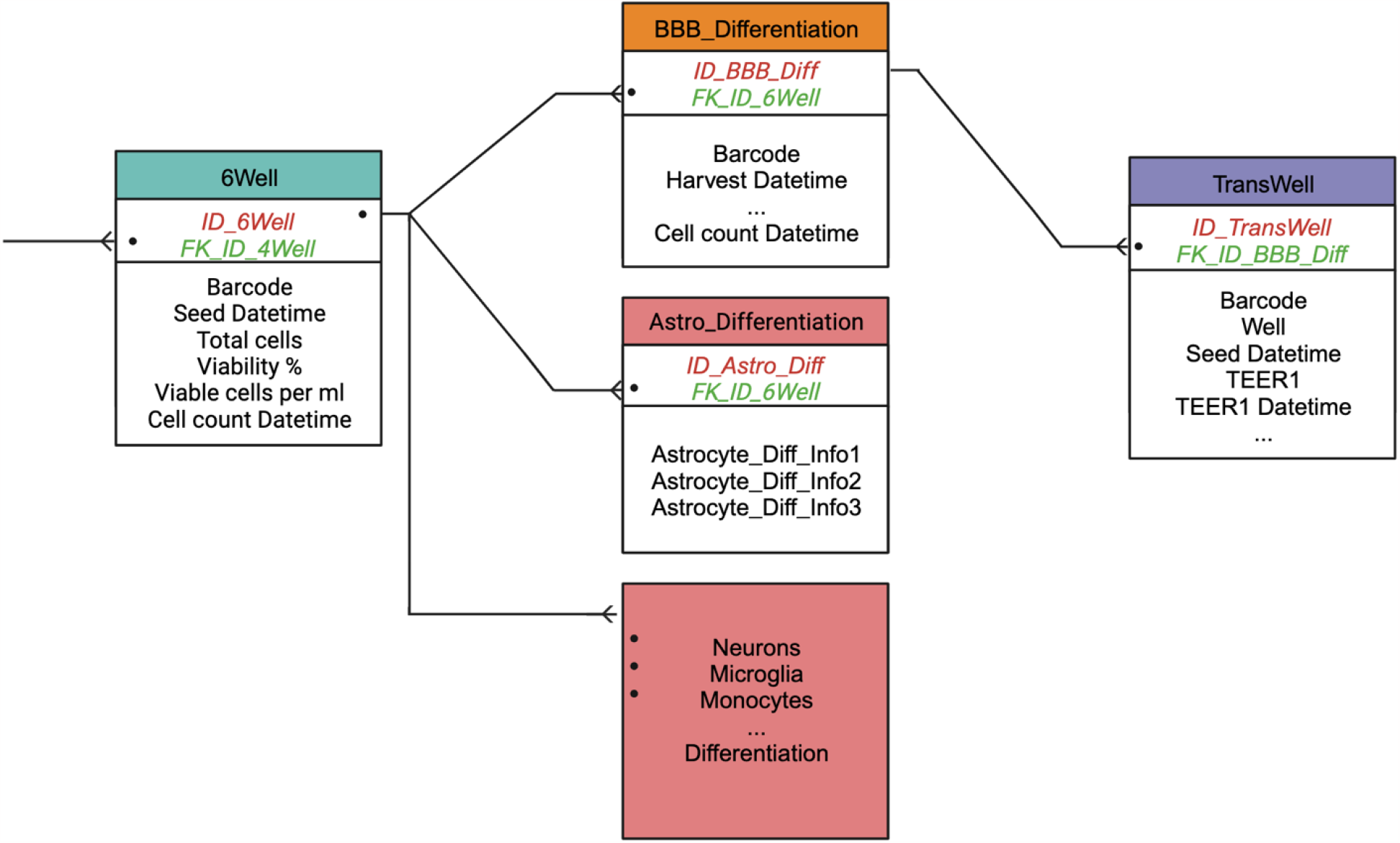
Conceptualizing ASCC database. Possible expansion of database structure towards differentiation of other cell types (such as astrocytes, neurons, microglia, monocytes; red tables). The structure is flexible and allows appending of tables to a 6-well plate table (green) that stores information on undifferentiated cells through a one-to-many relationship. In this way, different differentiation protocols can be accommodated in the database.

**Table S1**. Primers used for the amplification of heavy chains, as well as light Kappa (κ) and Lambda (λ) chains. Three PCR reactions are performed per chain. <u>PCR1</u> and <u>PCR2</u> are performed with forward and reverse primer mixes. <u>PCR3</u> is performed with specific primer pairs selected from 69 individual primers. The selection is based on sequencing results of the second PCR amplification and identification of used V(D)J alleles. See the attached Table S1.xlsx file.

## Footnotes

1 https://github.com/CRFS-BN/mABpy

2 https://github.com/CRFS-BN/mABpy/tree/main/simplified_pipeline

3 https://github.com/CRFS-BN/mABpy/tree/main/simplified_pipeline-adjusting-the-pipeline-to-your-own-needs

4 DOI: 10.5281/zenodo.8229164

5 DOI: 10.5281/zenodo.10106688xs

